# Measuring functional connectivity in stroke – approaches and considerations

**DOI:** 10.1101/177618

**Authors:** Joshua S. Siegel, Gordon L. Shulman, Maurizio Corbetta

## Abstract

Recent research has demonstrated the importance of global changes to the functional organization of brain network following stroke. Resting functional MRI (R-fMRI) is a non-invasive tool that enables the measurement of functional connectivity (FC) across the entire brain while placing minimal demands on the subject. For these reasons, it is a uniquely appealing tool for studying the distant effects of stroke. However, R-fMRI studies rely on a number of premises that cannot be assumed without careful validation in the context of stroke. Here, we describe strategies to identify and mitigate confounds specific to R-fMRI research in cerebrovascular disease. Five main topics are discussed: 1) achieving adequate co-registration of lesioned brains, 2) identifying and removing hemodynamic lags in resting BOLD, 3) identifying other vascular disruptions that affect the resting BOLD signal, 4) selecting an appropriate control cohort, and 5) acquiring sufficient fMRI data to reliably identify FC changes. For each topic, we provide evidence-based guidelines for steps to improve the interpretability and reproducibility of FC-stroke research. We include a table of confounds and approaches to identify and mitigate each. Our recommendations extend to any research using R-fMRI to study diseases that might alter cerebrovascular flow and dynamics or brain anatomy.

## Introduction

In stroke, a disruption to the brain’s vascular supply leads to infarction and structural damage (i.e. cell death) of gray/white matter. But stroke also produces remote changes in structurally normal brain areas by a variety of different mechanisms,^1^ as well as shifting of brain tissue. Remote changes have been reported in metabolism, cerebral blood flow, resting neural activity, and evoked neural response.^2^ Until recently, studying the relationship between distant functional disruption and cognitive deficits in humans was a nearly impossible task. In the late 1990-early 2000s human fMRI studies showed that functional abnormalities correlate with deficits in language,^3,4^ attention,^5^ and motor function^6^ and these abnormalities tend to normalize in parallel with recovery of function. These observations suggested that neuroimaging can identify altered neural function across many brain regions, and that this functional alteration may be valuable for understanding abnormal behavior. A limitation of the task fMRI approach to studying stroke is that patients must be able to complete the task in order to assess activation. Further, the use of multiple compensatory strategies during task performance can complicate the interpretation of task fMRI data.

Around this same time, the study of functional connectivity using resting functional magnetic resonance imaging (R-fMRI) was gaining momentum.^7^ R-fMRI measures intrinsic fluctuations in the blood oxygenation level dependent (BOLD) signal in the absence of a task. The correlation in these signals between brain areas is used to infer functional connectivity (FC). This approach had substantial appeal as a tool for studying stroke, offering a non-invasive and task-free paradigm for studying human brain connectivity at high spatial resolution.^8^ A study in 2007 found that hemispatial neglect was strongly correlated with disrupted connectivity in the dorsal attention system,^9^ suggesting the potential of R-fMRI for studying stroke. As evidence has mounted in the past decade, it has since become increasingly apparent that understanding behavioral deficits will require a complete description not only of lesion topography, but also of remote connectivity abnormalities.^9–17^ Resting functional MRI (R-fMRI) remains a promising tool to examine network level changes in stroke and recovery in humans.^8,18,19^

Interpretation of the correlation of BOLD fluctuations in healthy subjects frequently rests on numerous assumptions. For example, a critical assumption of most R-fMRI research is that neurovascular coupling is relatively consistent across brain areas, across time, and across individuals. Though imperfect even in a healthy population, such assumptions have enabled reliable mapping of spatial and temporal relationships between brain areas.^20^ When studying cerebrovascular disease, a normal hemodynamic response cannot be assumed.^21^ However, if certain additional steps are taken to empirically identify and control for relevant confounds, then measured FC-stroke relationships can be meaningfully interpreted.

The goal of this article is to provide evidence for the critical importance of these confounds, and explore best practices to manage them. We will first consider issues relating to inter-subject registration – focusing on volume-based registration errors caused by stroke and recommending surface-based methods to improve registration. We will then consider hemodynamic coupling – discussing evidence that vascular disease can produce changes in the magnitude and latency of the hemodynamic response, and recommending an approach to measure and remove this confound. Finally, we will consider selection of appropriate experimental controls and control subject selection. We cite published data where available but also provide additional analyses using data from a cohort of stroke patients studied at Washington University.^15^ This dataset is publicly available (see ‘Public Stroke Data’ below).

Table 1 summarizes stroke R-fMRI confounds and approaches to identify and mitigate each. Many of our recommendations are quite feasible in current FC-stroke datasets. Moreover, many of our recommendations are not specific to stroke only, but extend to any research using R-fMRI to study diseases that might alter cerebrovascular dynamics or brain anatomy.

**Table 1.**
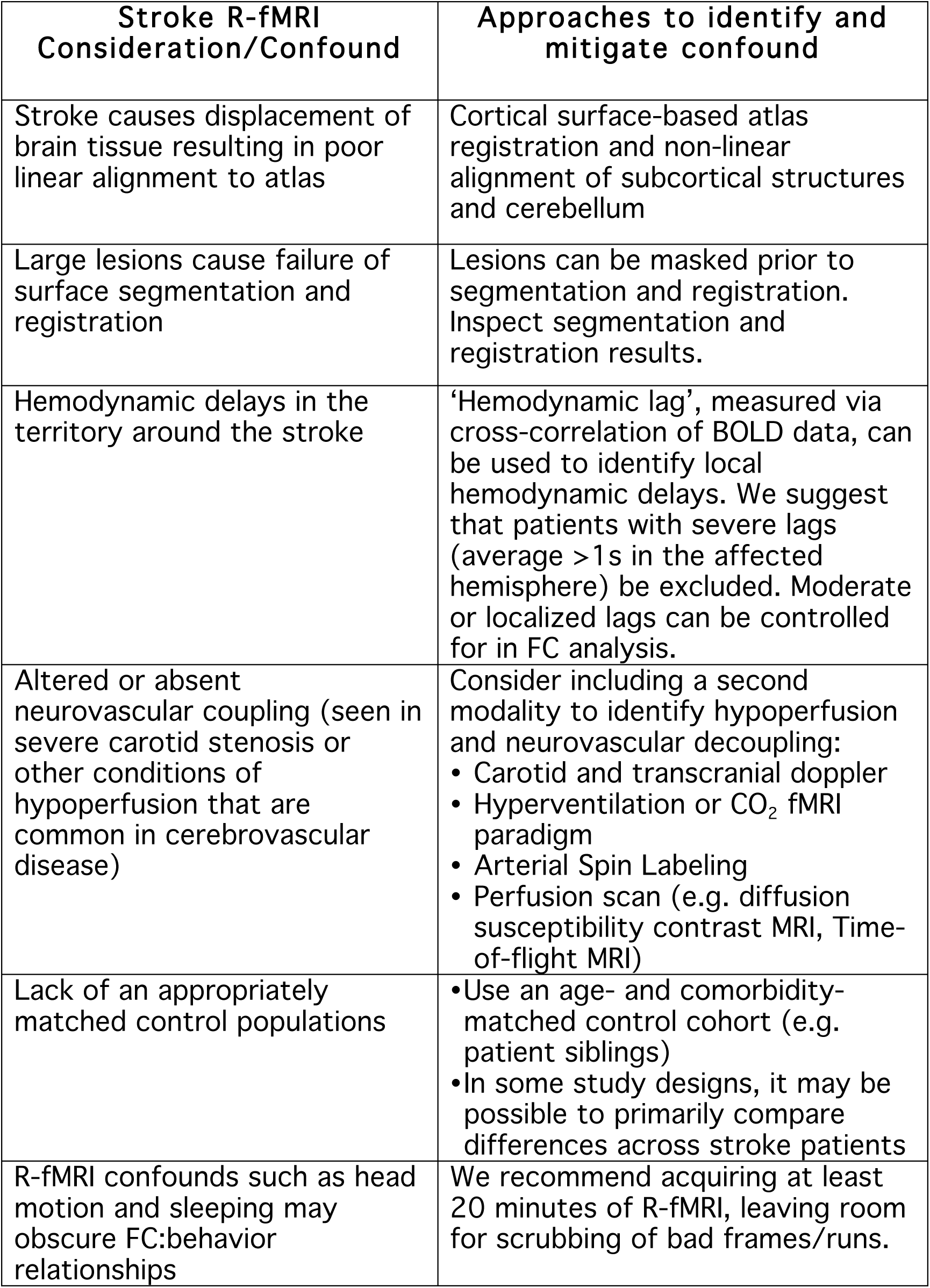

### Registration of the cortical surface and subcortex

Many previously published FC-stroke research has relied on registration to a common atlas space using 6-parameter affine linear transformation.^12,13,22,23^ The limitation with this approach is that a stroke, and pathophysiological processes associated with stroke can lead to substantial relative displacement of tissue (i.e. the central sulcus might move anterior or posterior relative to other brain landmarks). This ‘mass effect’ phenomenon is illustrated after volume alignment in data from 33 cortical stroke patients (Fig. 1). An 8mm radius sphere is placed in the angular gyrus - defined based on anatomical landmarks - in each individual linearly-aligned brain. A conjunction image shows good overlap in healthy individuals (with some voxels showing 100% overlap), but poor overlap in patients (with a maximum of 63% overlap). This was the case across the cortex. For 156 ROIs that span the cortex,^24^ ROI overlap was significantly lower in patients than controls (t = 9.6, p< 0.001). Non-linear alignment did not substantially improve seed co-localization in the angular gyrus. This may be because it uses only tissue contrast and not cortical folding patterns. Here, linear and nonlinear were compared using FSL (FMRIB Software Library) registration algorithms.^25^ A variety of linear and nonlinear brain registration software packages are available, so results may vary somewhat with different software. However, as discussed below, non-linear registration should substantially improve alignment of subcortical structures.

**Figure 1.**
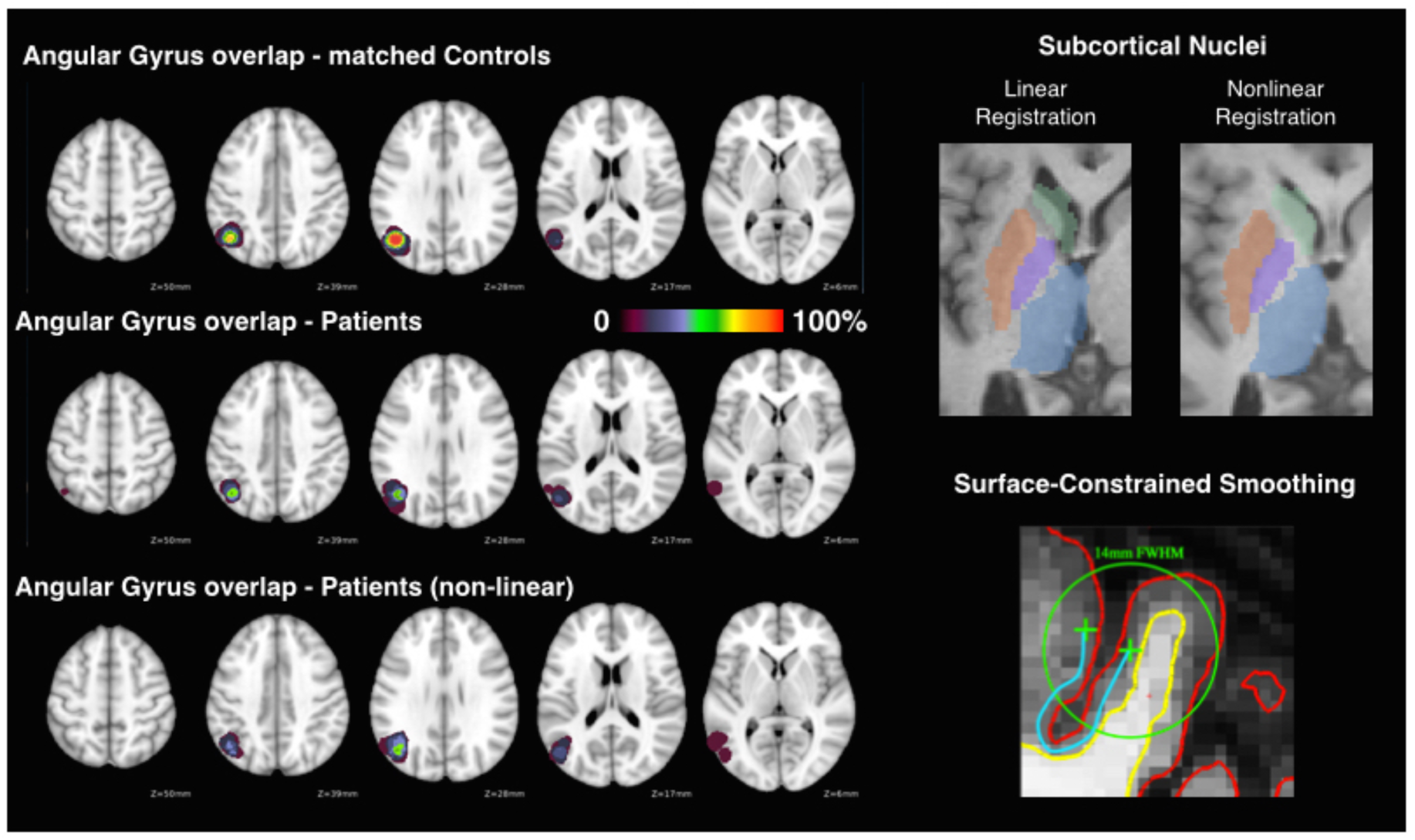
Nonlinear registration and surface-based tools improve stroke patient alignment. Quality of linear alignment is compared between 33 patients with large lesions (greater than 40 cm_3_) on either hemisphere and 24 matched controls. A region in the right central sulcus was defined in each subject following surface folding-based registration. Separately, each brain was linearly aligned to a reference atlas in Talairach space. The landmark-defined angular gyrus region was then projected to the volume coordinates in the linearly aligned brains. **Top right:** An example of registration of subcortical nuclei using linear versus nonlinear registration. Atlas-defined ROIs are shown for caudate, putamen, globus pallidus, and thalamus. Example patient (used in Figure 4) has a cortical lesion in the contralateral hemisphere. **Bottom Right**: A demonstration of surface smoothing enabled by FreeSurfer (Image courtesy of Dr. Douglas N. Greve). The green line indicates the full-width half-max boundaries of a 14mm volume-smoothing kernel. Notice that the smoothing kernel would cause functional data to be smoothed across multiple gyral walls. This problem is mitigated with surface smoothing (blue line).

Surface-based registration offers a solution to variability in cortical shape and variability in shifting of tissue after stroke.^26–30^ Surface- and contour-based alignment approaches provide superior registration. The benefits of surface-based registration have been demonstrated in the healthy brain^31–33^, and those benefits are magnified by the substantially increased heterogeneity resulting from stroke. However, to our knowledge, this is the first demonstration of the benefits of a surface-based approach in FC-stroke analysis.

For analysis of subcortical nuclei and the cerebellum, nonlinear volume alignment (such as FSL FNIRT) may also provide registration that is superior to linear registration (Fig. 1, top right). This is because patients’ brains often show anatomical shifts as well as enlarged ventricles. Even patients with more moderate lesions frequently show substantial enlargement of lateral ventricles, causing reduced quality of alignment of basal ganglia and thalamus.

Additional advantages exist to segmentation of tissue compartments beyond the important issue of registration. One is that surface segmentation enables surface-constrained smoothing, so that gray matter signal can be smoothed with minimal contamination by signal from CSF, white matter, or opposing gyral walls (Fig. 1, bottom right). Another advantage of segmentation is that it allows for high quality definition of tissue compartments in the individual that can then be used as nuisance regressors (white matter, CSF) in data cleaning approaches such as aCompCor.^34^

An important caveat of either nonlinear volume alignment or surface-based registration is that large lesions should be explicitly excluded or masked prior to alignment and the results should be carefully assessed for quality and feasibility. This is necessary in order to prevent the mislabeling of cortical structures.

Large lesions can cause distortion or failure of surface segmentation and registration. In such instances, we have found that painting over the lesion with voxel values from a T1-weighted brain atlas can improve landmark and folding based surface alignment approaches. This is demonstrated in Figure 2. In the top middle panel, the high contrast lesion (blue arrow) has caused massive disruption of the surface segmentation (yellow arrows). This causes errors in surface tessellation as well as parcellation (yellow dotted lines). But masking of the lesion enables FreeSurfer to properly trace the remaining cortical surface. The surface within the masked area can then be removed from analysis. An important consideration is that, while this fix has enabled segmentation to complete in all cases, the resulting segmentation will require visual inspection and possibly additional manual editing.

**Figure 2.**
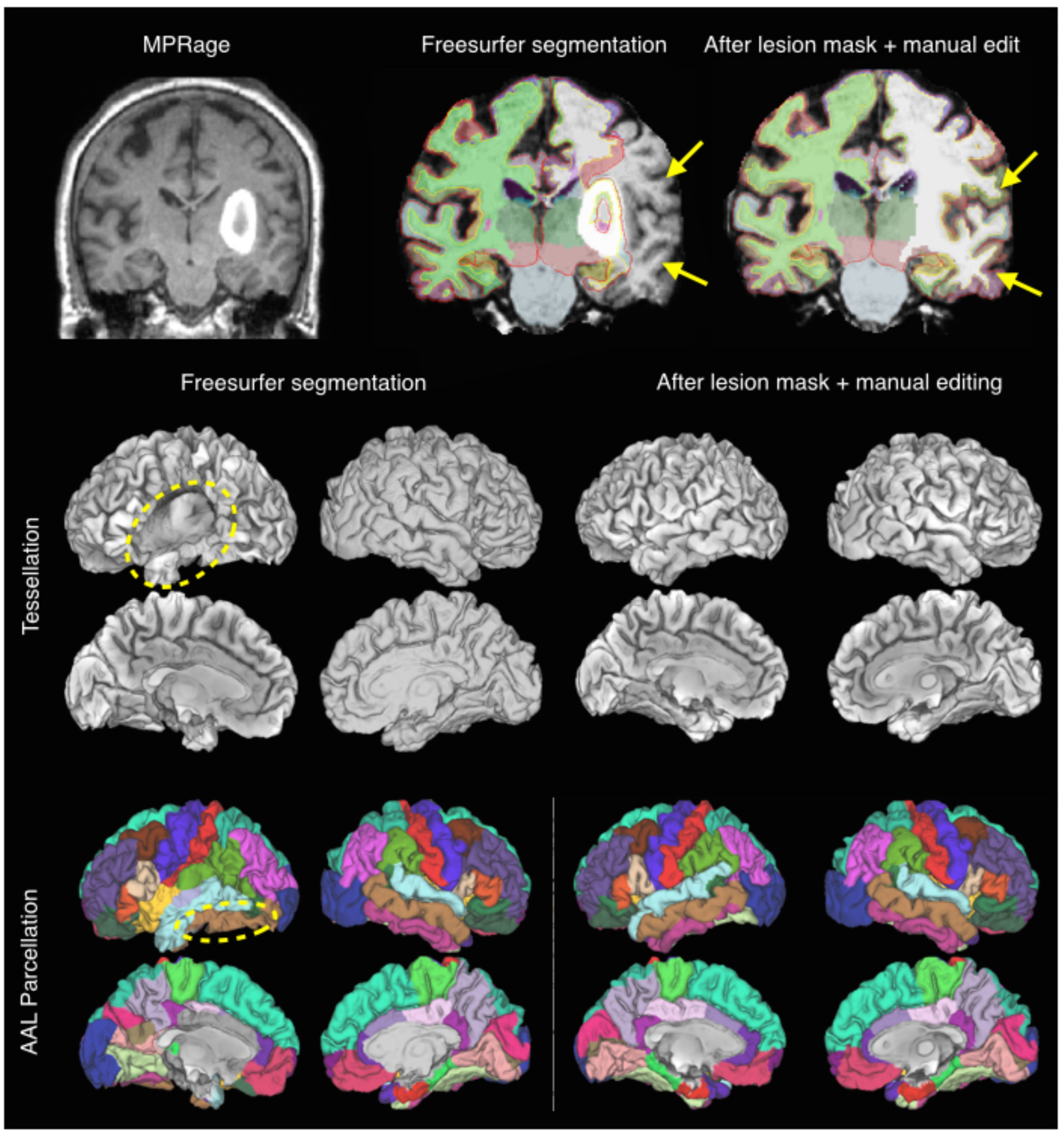
FreeSurfer segmentation error caused by a large lesion, and subsequent resolution after lesion-masking and manual editing. Top: MPRage illustrating a hyper-intense hemorrhagic stroke. Unattended FreeSurfer segmentation (middle) is unable to correctly identify the cortex lateral to the lesion. To resolve this, lesion masking with T1 atlas values and manual editing using control points is done. In the resulting segmentation (right), FreeSurfer correctly identifies and segments the cortical surface. **Middle and bottom:** The surface and AAL parcellation generated by the post-FreeSurfer HCP pipeline show errors in tracing of the cortical surface as well as labeling errors (indicated by yellow dotted line). After lesion masking and manual editing, these errors are no longer present.

An approach we have found for easily identifying errors in pial surface segmentation is viewing the pial surface segmentation on top of the MPRage. In Connectome Workbench, the pial surface can be color-coded based on FC values to further assess accuracy. Displaying homotopic FC values (which should always be positive) can aid in identifying segmentation errors (Fig. 3).

**Figure 3.**
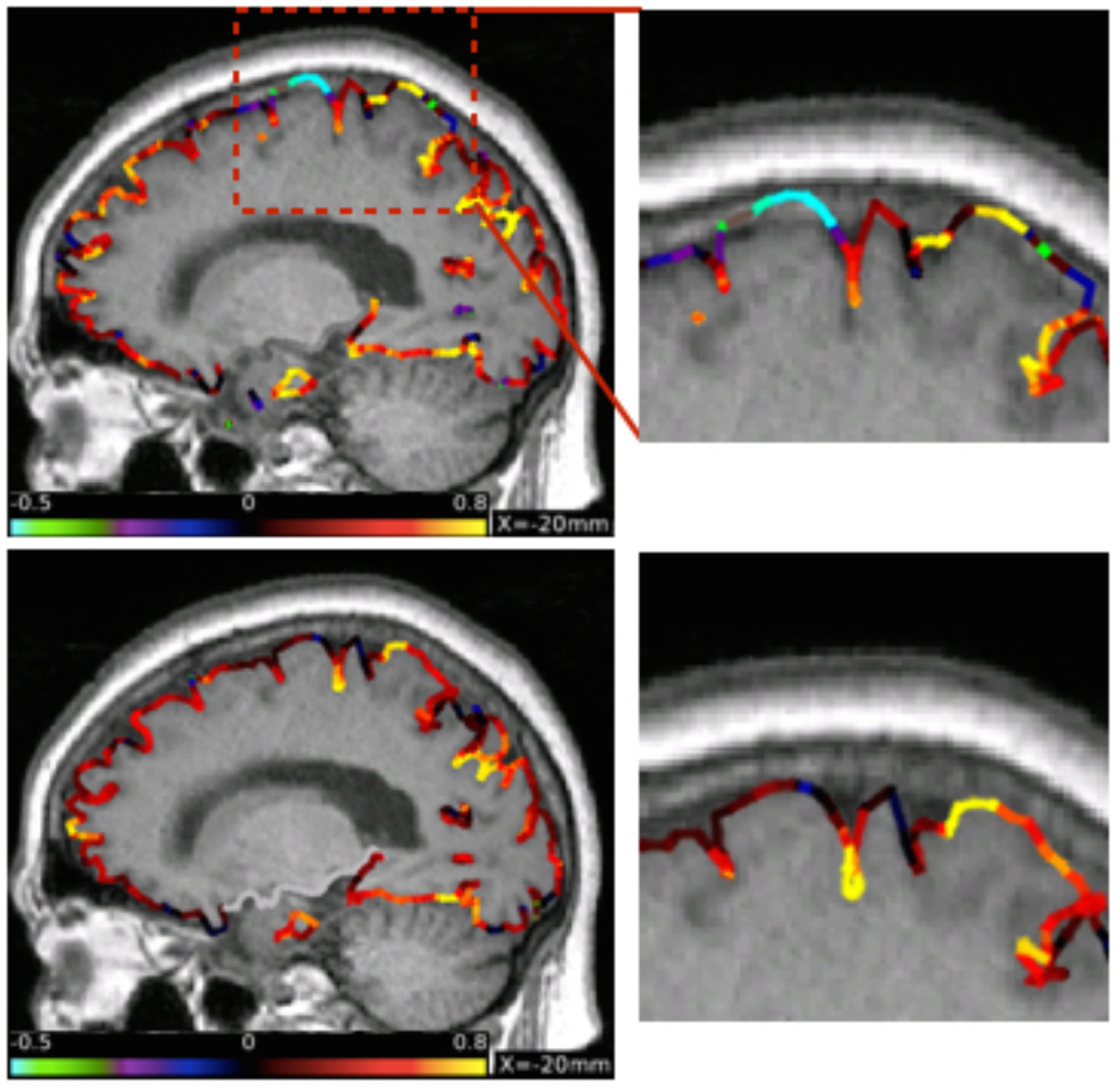
Identifying and correcting gray matter segmentation. Top: The FreeSurfer-defined pial surface is displayed as a ribbon over the subject’s MPRage. The ribbon is color-coded based on Homotopic FC strength at each surface vertex. In locations in which errors of inclusion of dura mater have occurred, low or negative homotopic FC is found. **Bottom**: After manual editing of FreeSurfer segmentation pail ribbon accuracy is improved and homotopic FC values are higher.

Following proper segmentation and surface registration, we use post-FreeSurfer preprocessing pipelines from the Human Connectome Project.^30^ Future improvements on the HCP pipeline may enable manual editing of FreeSurfer segmentation so that the entire HCP preprocessing pipeline is compatible with the approach described above.

Misalignment can a source of noise or confound in FC-stroke studies. As we have quantitatively and qualitatively demonstrated, after automated but manually optimized surface segmentation, and nonlinear alignment of subcortical nuclei, improved functional alignment is possible.

### Blood flow and hemodynamics

Task fMRI studies have shown reduced amplitude^21,35–37^ and increased latency after stroke^38–42^ - with BOLD responses sometimes peaking 15 seconds or more after transient neural activation. Changes in amplitude and latency have even been demonstrated in both affected and unaffected hemispheres and are most prominent in, though not limited to, the first three weeks after stroke.^35,37^ For a more extensive review of functional MRI studies in stroke, see.^43^ Importantly, changes in amplitude and latency often seem to co-occur, i.e. hemodynamically compromised vasculature shows a BOLD hemodynamic response function that is both reduced in amplitude and increased in latency.^39–41^

Fortunately, it is possible to identify latency changes. The remainder of the section discusses ways in which hemodynamic coupling is altered after stroke, means by which such alterations may be identified, and strategies to control for these confounds in FC-stroke analyses.

### Hemodynamic lags

In the last 3 years, a handful of reports have identified regional delays on the order of seconds in resting fMRI fluctuations in stroke and cerebrovascular disease patients.^44–49^ These delays are measured by cross-correlation (i.e. time shift analysis) of regional BOLD timecourses with some reference signal. This technique has been applied with different choices of reference signals including gray signal^47^, homotopic signal from the non-lesioned hemisphere^45^, and a superior sagittal sinus seed.^46^ Some benefits of the cross-correlation approach is that it is fairly robust to the choice of reference signal^46^ and that a hemispheric measure of lag severity can be attained from only six minutes of R-fMRI data^49^ – though greater spatial specificity requires longer scans.

Importantly, lag can be seen in areas of perfusion deficit as measured by contrast enhanced perfusion-weighted imaging^44,47^ and arterial spin labeling^49^ and also seems to occur in areas with reduced amplitude of evoked BOLD response.^45^ However, studies directly relating latency and amplitude of evoked response to lags in resting BOLD are still needed.

At two weeks post-stroke, the prevalence of patients showing substantial hemodynamic lags is above 30%. This number drops close to 15% by 3 months post-stroke and 10% by 1 year.^49^ Interestingly, lag severity correlates with lesion size as well as severity of deficits.^49^

As would be expected, lags systematically alter measurements of FC from the affected node. This is easily illustrated by comparing homotopic lags to homotopic FC. Homotopic FC provides a useful comparison because it is typically strong in the healthy brain, and because it has previously been associated with deficit after stroke.^9,16,50^ In figure 4 each circle represents a pair of ROIs in opposite hemispheres (homotopic) in one patient (red) or control (blue). The lag between homotopic ROIs is plotted on the x-axis while the functional connectivity (zero-lagged Pearson correlation) is plotted on the y-axis. This figure was generated using a cohort of 107 sub-acute stroke patients and 24 age-matched controls described in.^49^ This figure is particularly useful because one plot makes multiple important points – 1) hemodynamic lags produce a monotonic decrease in measured FC values, 2) even in areas with zero lag, homotopic FC is lower on average in patients than controls, 3) negative homotopic FC values observed in patients are likely (though not definitively) a symptom of lags, and 4) lags are common in patients and rare but reliably identified in a small minority of risk-matched controls (2/24 with consistent measures across scans 3 months apart).

**Figure 4.**
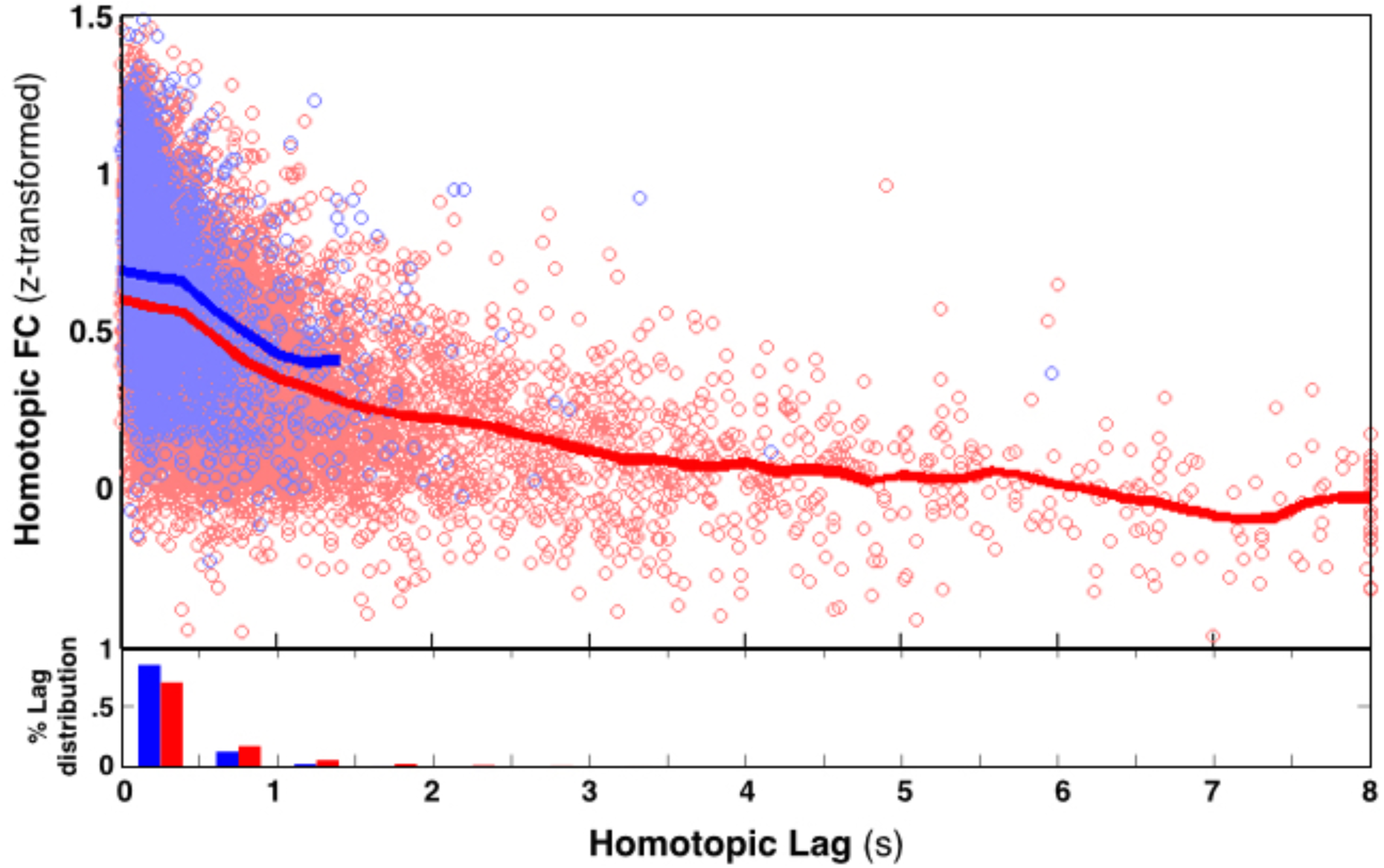
Hemodynamic lags systematically alter functional connectivity. Each circle represents a pair of the ROIs on opposite hemispheres (homotopic) in one patient (pale red) or control (pale blue). The lag between homotopic ROIs is plotted on the x-axis while the functional connectivity (zero-lagged Pearson correlation) is plotted on the y-axis. The LOWESS moving average is plotted in bright red and blue lines for patients and controls, respectively. This figure is generated using the 107 sub-acute stroke cohort and 24 age-matched controls. A set of 78 left hemisphere ROIs (and their right hemisphere mirror image regions) were used for each subject. ROIs intersecting a lesion were excluded. Under, a histogram shows the proportion of homotopic ROI pairs showing lags. Even in sub-acute patients, the majority of regions show a lag of less than 0.5 seconds.

Interestingly, homotopic anti-correlations appear in many FC-stroke studies,^9,51^ suggesting that the studies were likely affected by lag. A few approaches have been proposed to correct for lags in FC analysis. Bauer et al., proposed that using the timecourse from voxels within the lesion as a nuisance regressor reduced aberrant observations attributable to lag in mice with transient MCA occlusions. Another approach that has been suggested is to shift timecourses in lagged tissue (backwards in time to ‘reverse’ measured lags) prior to FC analysis.^46,49^ We do not endorse either approach for the following reasons. Areas of lag also show reduced BOLD signal power in much of the infra-slow FC range (.045-0.09Hz).^49^ In affected tissue, two changes occur: 1) a temporal delay, and 2) a change in frequency content. We would expect that temporally shifting timecourses might correct the first change, but will not correct the second change. This may explain the observation that ‘correcting’ timecourses reduces FC aberrancy, but does not remove it (see Siegel et al., 2015, figure 5). There is no practical way to correct a change in the frequency content of the BOLD signal. Moreover, because other sources of unwanted variance (head motion, white matter, CSF signals) may or may not be shifted in lag-affected areas, shifting timecourses might disrupt nuisance regression in undesirable ways.

**Figure 5.**
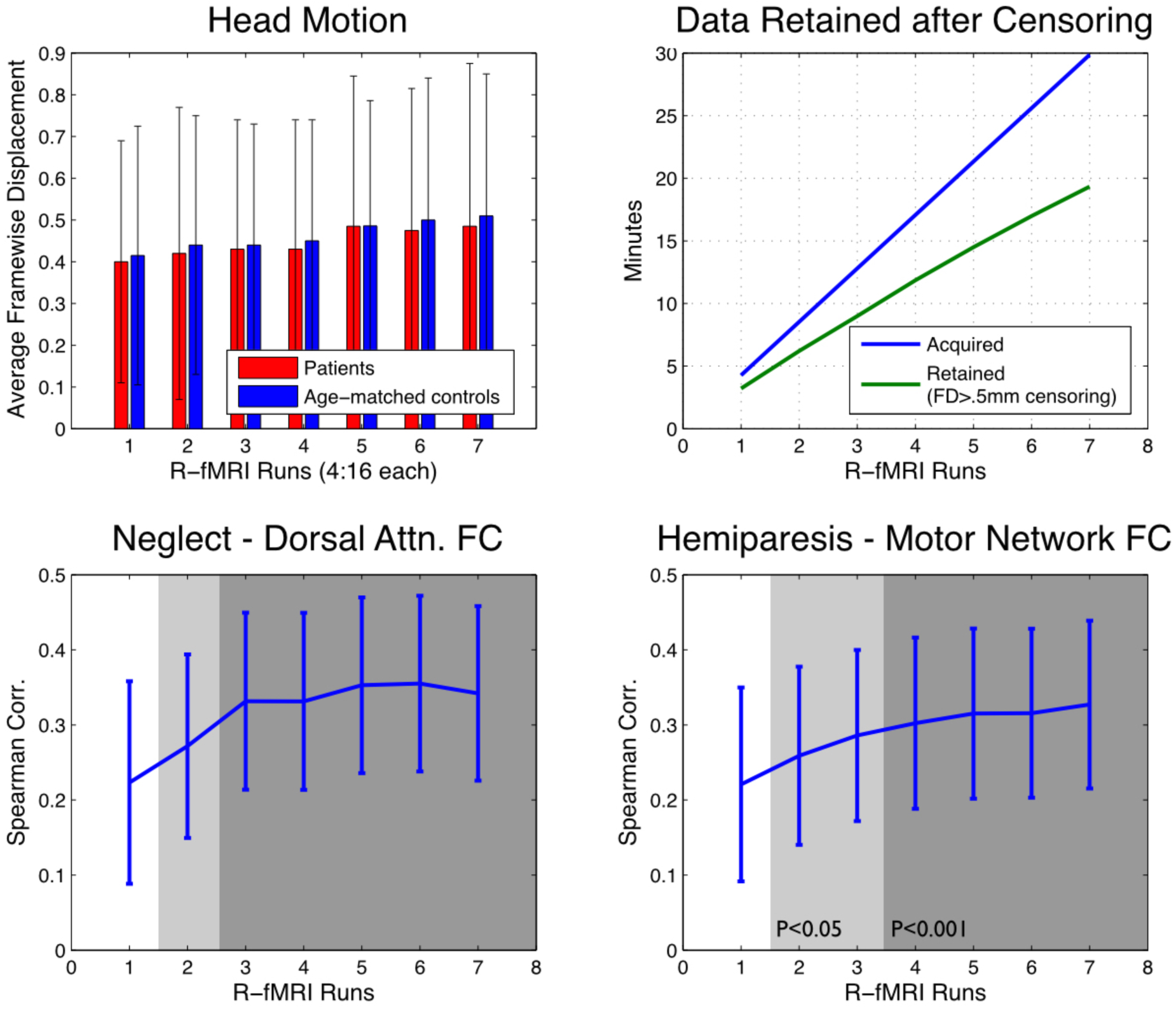
More R-fMRI data per subject improves FC-Behavior Correlation. To assess the relationships between scan length and FC-behavior correlation, we generated FC estimates using fractions (1-7 individual 4:16 runs) of our 7 run session in N=96 stroke subjects. In each sub-sample, two canonical FC-behavior relationships were estimated across the sub-acute stroke cohort. **Top Left:** Average head motion (framewise displacement) during each of seven consecutive runs. **Top Right:** The blue line shows minutes of data acquired, the green line shows average number of usable frames after exclusion of frames with framewise displacement > 0.5mm. **Bottom Left:** visual field bias (the deficit of which produces hemi-spatial neglect) was correlated with interhemispheric FC in the dorsal attention network. **Bottom Right:** Contralesional motor function is correlated with inter-hemispheric FC in the hand/body somatomotor network. Error bars indicate 95% confidence interval of Spearman’s rho between FC:behavior.

Instead, we recommend that the most extreme lag cases be excluded from further analysis and smaller changes be considered after FC is calculated. In subjects with severe and widespread lags (for example, average homotopic lag >1second i.e. the entire lesioned hemisphere is delayed by greater than 1 second relative to the contralesional hemisphere) FC data will be so altered as to be unusable in analysis. In our opinion, exclusion of such subjects is the only feasible option. While 1 second is arbitrary, it is a conservative cutoff that should only exclude roughly 5% of ischemic stroke subjects (6/119 patients at 2 weeks in our dataset). In patients with a constrained area of large magnitude lag (as is seen in the example patient in Fig. 4 – area of lag is highlighted with a green outline) it may be beneficial to exclude the affected region from FC analysis. Since the effects of lags of 1-6 seconds are approximately linear (see Fig 5 of Siegel et al., 2015), more subtle lags can be co-varied out of any FC analysis.^52,53^ Fortunately, this precaution is feasible because lags can be calculated using the same data used for canonical FC analysis. Researcher investigating FC-stroke data can test for lags using publicly available software (nil.wustl.edu/Corbetta/resources/lagsuite.tar.gz). Regardless of the approach taken, it is important for investigators to bear in mind that aberrant FC in cerebrovascular disease patients may reflect both vascular and neuronal changes.

### Neurovascular Coupling

While identifying lags is important, it may not be a complete fix for the challenge of altered neurovascular coupling. In some patients, a decrease in amplitude^35,54^, or a complete loss of the BOLD response^37,55^ has been observed in the absence of lags. On it’s own, it is difficult to interpret reduction/absence of a evoked BOLD response – it might reflect a decrement in neural activity, loss of neurovascular coupling, or an inability of the vasculature itself to adequately increase local perfusion.^36,37,56^(Marshall, 2004; Pineiro et al., 2002; Rossini et al., 2004) Thus, approaches to better validate hemodynamic responsiveness to neural activity would be of value.

Carotid stenosis is perhaps a useful example of a vascular disease because it is relatively well characterized in its effects on neurovascular coupling, it is common in stroke patients, and it’s effects on FC have been reported. Severe carotid stenosis (>70% occlusion) reduces both the static and dynamic components of cerebral blood flow.^57^ This results in uncoupling of the hemodynamic response from neural activity.^58^ Moderate stenosis (50-70% occlusion) may not alter coupling,^59^ though this has not been carefully addressed. In a healthy asymptomatic population of adults over the age of 70, 4.8% exhibit moderate (>50% occlusion) and 1.6% exhibit severe (>70% occlusion) carotid stenosis.^60^ In a stroke population, prevalence is substantially higher. Based on clinically acquired carotid doppler data from our stroke cohort, 20% of patients (13/66) show >50% occlusion with 11% (7/66) showing >80% occlusion on the affected side. Other studies have reported that as much as 33% of ischemic stroke patients exhibit moderate or severe intracranial stenosis, and 12% have stenosis on the hemisphere opposite the lesion ^61^. In most such cases, an embolic stroke produces an infarct in only a portion of the territory affected by the stenosis. Studies directly measuring FC changes in carotid stenosis patients relative to non-stenotic controls have explicitly shown large reduction in FC in the affected hemisphere ^62,63^.

Though it is intuitive that changes in neurovascular coupling should alter FC measurement, the effects are not straightforward. Relatively small decreases in the magnitude of the HDR might have little effects because functional connectivity analysis typically relies on correlation (not covariance) of the r-fMRI signal. However, it is also possible that changes to neurovascular coupling can alter FC in profound ways. Further studies are required to understand the relationships between abnormalities of neurovascular coupling and FC. One goal of such studies would be to develop better techniques to identify and control for vascular changes in FC analysis. This is possible by employing a combination of R-fMRI with modalities for vascular imaging and electrophysiology. A challenge in such studies will be the fact that relationships between cerebral perfusion, cerebral autoregulation, and cerebrovascular coupling in the context of ischemia are exceptionally complex (for a review, see ^64^).

Some measures can be taken to identify pertinent vascular disruptions in FC-stroke analysis. These include assessment of internal carotid stenosis using carotid doppler, or assessment of local neurovascular coupling using CO_2_ or hyperventilation fMRI paradigms.^35,65^ Other imaging techniques capable of identifying clinically silent carotid stenosis include angiography (by MR and CT), transcranial doppler (which can be used in combination with arterial blood pressure to assess auto-regulatory impairment),^66^ and possibly measures of static regional cerebral blood flow (though tissue at the edge of the autoregulatory range may show normal rCBF but no hemodynamic response).

Pulsed arterial spin labeling (ASL) techniques are frequently used as a non-invasive measure of perfusion, though decreased signal cannot be interpreted as rCBF change as it may also reflect changes in mean transit time^67,68^ and may not be effective at identifying perfusion deficit in stroke.^69^ However, advances in ASL are making it possible to classify multiple features of vascular flow using multi-delay sequences to more accurately model cerebral blood flow even in circumstances of altered transit time.^23^

### General Recommendations

Stroke patients as a population have a significantly elevated prevalence of numerous medical comorbidities. These include diabetes, hypertension, cardiovascular disease, cerebrovascular disease (such as carotid stenosis), white matter disease, and others.^70^ For these reasons, we give three additional recommendations for studies applying R-fMRI in stroke patients.

#### 1. Choosing an appropriate control population

While it may be difficult to perfectly match controls on all relevant health factors, an approach that can substantially reduce the influence of such factors is to use siblings of patients as controls.^71^ An alternative approach that may be equally valid for some experiments is to compare performance measures across heterogeneous stroke patients (i.e. compare patients with and without a particular deficit). This allays the challenge of substantial heterogeneity in any human stroke population.

#### 2. Acquiring sufficient data

In healthy subjects, the reliability of functional connectivity measurements increases rapidly up to about 12-15 minutes of good data,^24,72,73^ while high-specificity single subject FC measurement or more complex parcellation approaches require even longer scan times, with reliability improving steadily up to 30 minutes.^74^ In-scanner head motion and propensity to fall asleep are not significantly different between stroke patients and age-matched controls (unpublished data), however in all data it is important to exclude corrupted scans.^75^ Notice that both groups increased in head motion somewhat with successive scans (Fig. 5, top left). Shorter runs with breaks and opportunities to stretch in between may help to mitigate deterioration in data quality.

After censoring frames corrupted by motion (framewise displacement > 0.5),^76^ we found that approximately 2/3 of fMRI frames were usable (Fig. 5, top right). Thus, we recommend acquiring at least 20 minutes of R-fMRI for each stroke patient, as that this is necessary to provide 12-15 minutes of good data.

To assess the relationships between scan length and stroke FC:behavior relationships, we generated FC estimates on N=96 stroke patients using fractions (1-7 individual 4:16 runs) of our 7 run session protocol. We compared these FC estimates to behavior using canonical (previously published and replicated) relationships – 1) hemispatial neglect and reduced interhemispheric FC in the dorsal attention network, and 2) hemiparesis and reduced interhemispheric FC in the somatomotor network. Using these sub-samples of the data, we found that canonical FC:behavior relationships pass significance (p<0.05) when only two sessions (∼8.5min of BOLD data) are used, but the relationships continue to become less noisy as longer scans are included (Fig. 5, bottom). Exactly how much R-fMRI data is necessary in stroke patients will depend on the goals of the study.

For these analyses, we used the following data inclusion/exclusion criteria - any frame with FD > 0.5mm was censored, and any run in which >50% of frames were censored was excluded entirely, and only patients with at least 3 minutes (90 frames) of data were included for FC analysis. Out of a total of 96 subjects with imaging and behavioral data, the number patients included in FC:behavior correlation was: N_1_=63 [i.e. number of usable subjects using only the first acquired run = 63], N_2_=74, N_3_=80, N_4_=80, N_5_=81, N_6_=82, N_7_=83. The increase in number of usable subject also contributed to improving p-values for FC:Behavior correlations.

#### 3. Manual checks on processing steps

We emphasize that anatomical and pathophysiological changes specific to stroke introduce increased possibility for errors in seemingly trivial processing steps that normally proceed without error in healthy subjects. Automated processing pipelines may allow such errors to go unnoticed. We recommend separate manual assessment of segmentation, preprocessing, and FC processing. As an example of how this can be done, we provide a set of images that we check in our patients to assess data quality (Fig. 6). Images on the left are primarily useful for checking surface segmentation. As shown in the right middle panel, we inspect a lag map for every subject. In some cases, a spatially constrained area in the vicinity of the lesion shows severe lags (green outline). This can be masked and excluded from FC analysis. As described previously, visualizing surface homotopic FC on top of an average volumes image can serve as an additional way to assess registration of functional and structural data.

**Figure 6.**
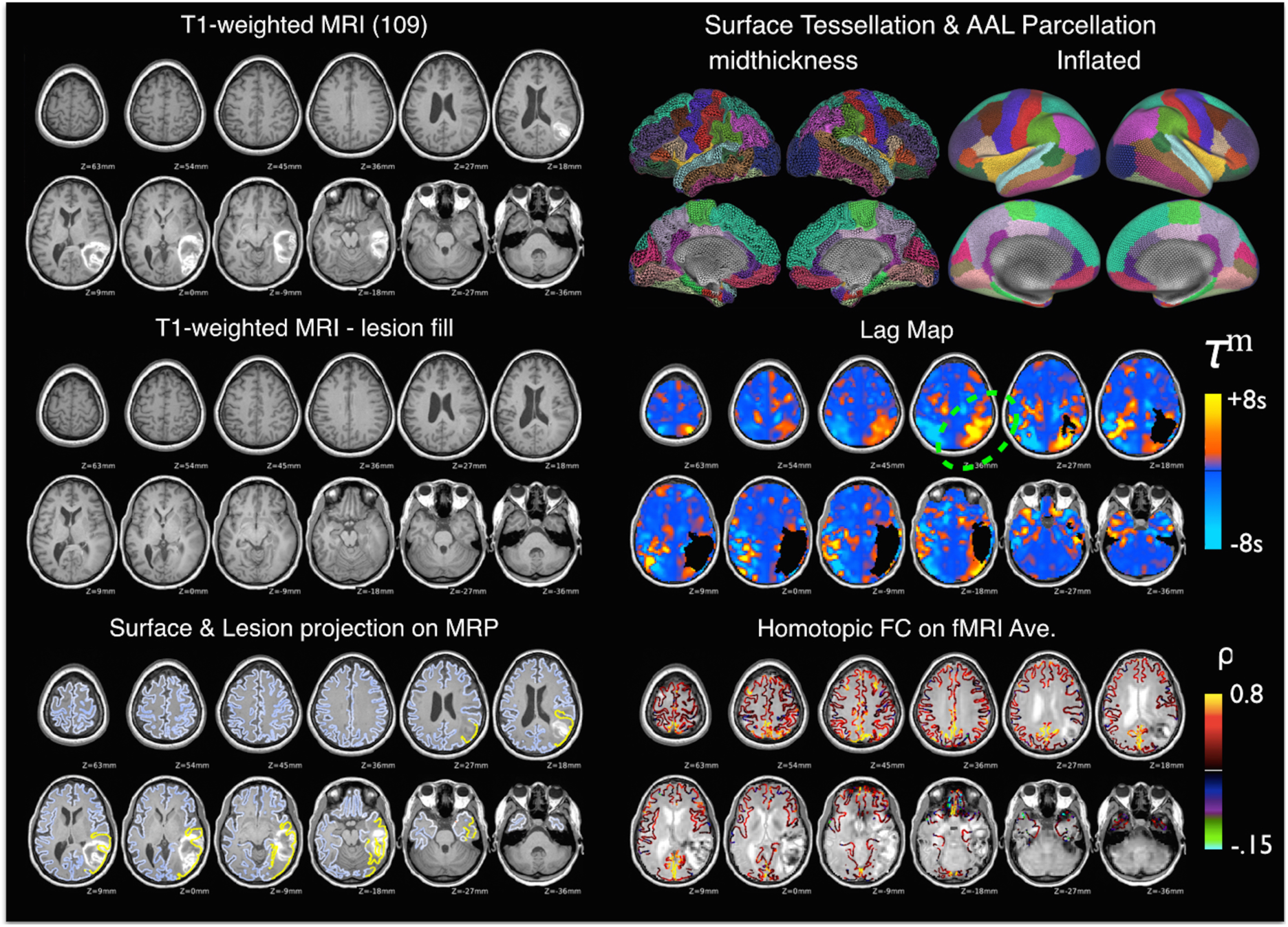
Quality assessment images for a sample subject. Top Left: T1-weighted MRI. For this subject, FreeSurfer segmentation failed due to the large hyperintense hemorrhagic stroke. The manually identified lesion was temporarily painted over with a T1-weighted atlas (**Middle Left**), allowing brain segmentation to run to completion. **Bottom Left:** cortical surface ribbon displayed on top of the T1-weighted MRI. Yellow denotes vertices included in a surface lesion mask. This is particularly useful to determine proper labeling of lesion and exclusion of dura in brain segmentaiton. **Top Right**: results of FreeSurfer segmentation - this can be compared to reference AAL parcellation to ensure proper labeling of structural features such as superior temporal sulcus, superior frontal sulcus. **Middle Right:** Lag map using homotopic reference (code available at nil.wustl.edu/labs/corbetta/resources). Areas of substantial lags are identified dorsal to the lesion boundaries and will be corrected or excluded in FC analysis. **Bottom Right:** homotopic FC overlaid on an average of aligned functional MRI volumes. Useful for checking alignment of functional volumes with MPR and surface segmentation.

We have mainly limited Figure 6 to images especially useful for identifying stroke- and comorbidity-related problems. We also use approaches to assess head motion, artifact, and BOLD signal quality that are commonly used in FC studies of normal populations, but discussion of those issues are beyond the focus of this paper (for a good discussion of those measures, see ^76^).

### The validity and value of functional connectivity stroke research

There are numerous reasons why prior FC-stroke findings represent promising advances, and why further FC-stroke research is merited. Post-stroke FC changes, such as reduced homotopic connectivity, are present across species and after careful consideration of confounds such as hemodynamic lags,^45^ are corroborated with other modalities that avoid neurovascular issues (such as EEG,^77^ voltage sensitive dye^78^ and axonal tracers^16^). Moreover, reported correlations between FC and behavioral deficits have proven robust and reproducible. Hemiparesis and hemispatial neglect provide useful examples. Hemiparesis is reproducibly associated with deficit in interhemispheric motor FC^14–16,50^ and recovery from hemiparesis parallels return of interhemispheric motor FC.^13,51,79^ Hemispatial neglect is reproducibly associated with reduced interhemispheric FC in attention networks^9,50^ and recovery from neglect parallels return of attention network FC.^53^ These two FC-behavior relationships can be doubly dissociated within a stroke population,^52^ and patients with similar deficits but heterogeneous lesions tend to show common patterns of FC disruption.^10,14,15^, suggesting that functional brain imaging provides important information beyond that of structural imaging.

FC stroke studies are particularly valuable for understanding the network-wide effects of stroke, because they provide the simultaneous assessment of multiple networks. It remains an open issue if acute FC studies can provide information about long-term outcomes, or whether they can be used to track the efficacy of therapy. Therefore much work lies ahead. This review highlights some of the potential methodological pitfalls of this approach and some of the ways to avoid or minimize these confounds. We hope that these recommendation will be helpful to the community not only of researchers in stroke, but any other pathology (eg. Tumor, neurodegenerative conditions, trauma, etc.) in which structural or neurovascular factors can affect the indirect but reliable relationship between neural activity and fMRI signals.

### Public Stroke Data

We have publicly released a stroke functional neuroimaging dataset (n=132 patients, n=31 age-matched controls) on the Central Neuroimaging Data Archives (available through https://cnda.wustl.edu/app/template/Login.vm - Study ID: CCIR_00299). This dataset includes structural imaging, functional MRI, neuropsychological testing scores across a wide range of behavioral assessments, demographics, arterial spin labeling (collected in a subset of patients) and carotid doppler ultrasonography classification acquired at two weeks post-stroke. Study design is described in its entirety in Corbetta *et al*., 2015.^71^ All data shown above is from that dataset. Figures 2,3 and 5 were generated by our functional connectivity processing pipeline available at www.nil.wustl.edu/labs/corbetta/resources/ and visualized using Connectome Workbench.

## Acknowledgement

This work was supported by the National Institute of Child Health and Human Development R01 HD061117-05 and the National Institute of Neurological Disorders and Stroke, R01 NS095741 (M.C.), and the American Heart Association, 14PRE19610010 (J.S.S.).

## Author Contribution

J.S.S., G.L.S. and M.C. all participated in writing and revising this manuscript.

